# Pleiotropy in enhancer function is encoded through diverse genetic architectures

**DOI:** 10.1101/188532

**Authors:** Ella Preger-Ben Noon, Gonzalo Sabarís, Daniela Ortiz, Jonathan Sager, Anna Liebowitz, David L. Stern, Nicolas Frankel

**Author notes:** These authors contributed equally to this work (E.P.B.N. and G.S. are listed in alphabetical order). Corresponding authors: David L. Stern < > and Nicolas Frankel.

## Abstract

Developmental genes can have complex c/s-regulatory regions, with multiple enhancers scattered across stretches of DNA spanning tens or hundreds of kilobases. Early work revealed remarkable modularity of enhancers, where distinct regions of DNA, bound by combinations of transcription factors, drive gene expression in defined spatio-temporal domains. Nevertheless, a few reports have shown that enhancer function may be required in multiple developmental stages, implying that regulatory elements can be pleiotropic. In these cases, it is not clear whether the pleiotropic enhancers employ the same transcription factor binding sites to drive expression at multiple developmental stages or whether enhancers function as chromatin scaffolds, where independent sets of transcription factor binding sites act at different stages. In this work we have studied the activity of the enhancers of the *shavenbaby* gene throughout D. *melanogaster* development. We found that all seven *shavenbaby* enhancers drive gene expression in multiple tissues and developmental stages at varying levels of redundancy. We have explored how this pleiotropy is encoded in two of these enhancers. In one enhancer, the same transcription factor binding sites contribute to embryonic and pupal expression, whereas for a second enhancer, these roles are largely encoded by distinct transcription factor binding sites. Our data suggest that enhancer pleiotropy might be a common feature of c/s-regulatory regions of developmental genes and that this pleiotropy can be encoded through multiple genetic architectures.

## Introduction

Developmental genes can have complex c/s-regulatory regions, with multiple enhancers scattered across stretches of DNA spanning tens or hundreds of kilobases [1-4]. Over many years, numerous studies have revealed a remarkable modularity of enhancer function, where distinct regions of DNA, bound by combinations of transcription factors, drive gene expression in defined spatio-temporal domains [5]. It has long been hypothesized that enhancer modularity facilitates evolution, because mutations in one enhancer can alter gene function without affecting the activity of other enhancers, thereby minimizing pleiotropic effects [6-8]. It is not clear, however, if the apparent modularity of enhancers reflects ascertainment bias, since few studies have looked explicitly for enhancer pleiotropy.

Many of the genes that regulate development have a pleiotropic role and their function is required in multiple developmental stages. A paradigmatic case of pleiotropy is that of *Hox* genes, a family of master transcription factors that specify the identity of body parts [9]. Recently, it has been uncovered that the same mechanism activates *Hox* genes in different organs of the mouse: the same enhancers activate *Hox* genes in both digits and genitalia [10]. Clearly, this implies that enhancers can have pleiotropic functions. It is not evident, however, whether these enhancers employ the same transcription factor binding sites to drive expression at multiple developmental stages or whether enhancers function as chromatin scaffolds, where independent sets of binding sites act at different stages.

*shavenbaby {svb)* encodes a transcription factor that orchestrates the differentiation of non-sensory cuticular projections (hereafter called trichomes) in *Drosophila melanogaster* [11,12]. *Svb* expression has been studied in detail mainly in the late embryonic stages, where it directs development of the epidermis and, concomitantly, the first-instar larval cuticle ([13], Fig 1B). *Svb* is also expressed in the pupal epidermis where it is required for trichome development in part of the wing, notum and abdomen [14,15] and for proper development of leg joints [16].

**Fig 1.**
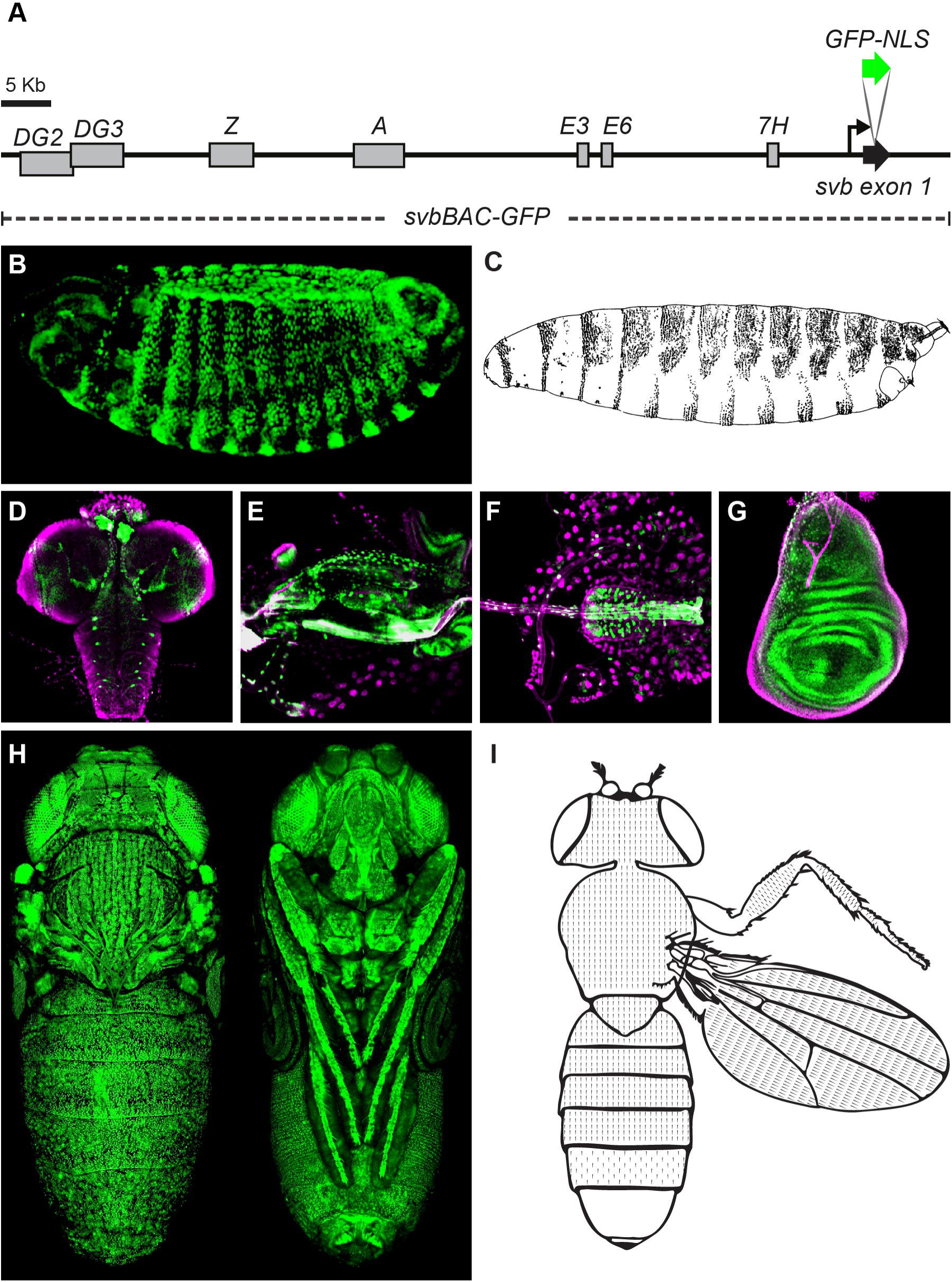
*svb* expression throughout *Drosophila melanogaster* development. (A) Schematic representation of svbBAC-GFP. Gray boxes represent the seven embryonic enhancers. The site of insertion of the GFP-NLS is indicated in the scheme. (B) GFP expression recapitulates the expression pattern of *svb* in the embryo. (C) Trichome pattern of the first-instar larva. (D-G) GFP expression in non-epidermal structures of the third-instar larva: central nervous system (D), pharynx and salivary glands (E), esophagus and proventriculus (F), and wing imaginal disc (G) DAPI stain in magenta. (H) GFP expression in pupal epidermis. (I) Representation of the trichome pattern in the dorsum of an adult fly.

The c/’s-regulatory region of the *svb* gene has been experimentally dissected in D. *melanogaster* [17-21]. We have shown that the embryonic expression of *svb* is generated by seven enhancers that are located in a −80 kb region upstream of the transcription start site of the gene ([11], Fig 1A). These seven enhancers drive partially overlapping expression patterns in the late embryo, and these overlapping patterns are required for robust gene expression [20]. Evolutionary changes in five of these enhancers led to reduced *svb* expression in the dorsum of the *D. sechellia* embryo, resulting in differentiation of naked cuticle, rather than trichomes, in *D. sechellia* [11,20,21].

In this work we show that all seven *svb* enhancers drive gene expression in multiple tissues and developmental stages at varying levels of redundancy. We have explored how this pleiotropy is encoded in two of these enhancers. In one enhancer, the same transcription factor binding sites contribute to embryonic and pupal expression, whereas for a second enhancer these roles are largely encoded by distinct sites. Our data suggest that enhancer pleiotropy might be a common feature of c/’s-regulatory regions of developmental genes, and that this pleiotropy can be encoded through multiple genetic architectures.

## Results

### Shavenbaby is expressed in the larval and pupal epidermis

To characterize the expression of *svb* in larval and pupal tissues, we engineered a BAC carrying the complete c/s-regulatory region of *svb* by placing the coding sequence of a nuclear GFP upstream of svb ATG (Fig 1A). We stably integrated this BAC, named svfc>BAC-GFP, in the fly genome through attP/attB recombination. We confirmed that svbBAC-GFP recapitulates expression of the native gene in embryos (Fig 1B). This epidermal expression prefigures the location of trichomes in the first-instar larva cuticle (Fig 1C). We then examined *svb* expression in later stages. We observed GFP expression in the epidermis of third-instar larvae (data not shown). This may reflect persistence of the GFP reporter from second-instar larvae, when *svb* expression is probably required to cause differentiation of trichomes that will decorate the cuticle of third-instar larvae. We also detected GFP expression in larval non-epidermal structures of ectodermal origin that do not produce trichomes. Specifically, we observed GFP in the central nervous system (Fig 1D), the foregut (Figs 1E-F), the imaginal discs (Fig 1G), and the trachea (data not shown). We also found that svbBAC-GFP pupae display GFP expression in all epidermal tissue (Fig 1H), which is consistent with the fact that the adult exoskeleton is almost completely covered with trichomes (Fig 1I).

### Shavenbaby is required for the formation of many, but not all, adult trichomes

Next we asked whether *svb* function is required for trichome development in the pupal epidermis, as it is in the embryo. Flies carrying *svb* null mutations normally die before they eclose, but we identified a few male escapers (carrying a null *svb* allele on their single X chromosome) that allowed us to assess the requirement of svb for trichome development in the adult cuticle. Escapers had fewer trichomes in the wing, the legs, and the dorsal abdomen than control flies, but they still retained trichomes over much of the exoskeleton (Fig 2). However, in most regions, trichomes were smaller than normal or were misshapen.

**Fig 2.**
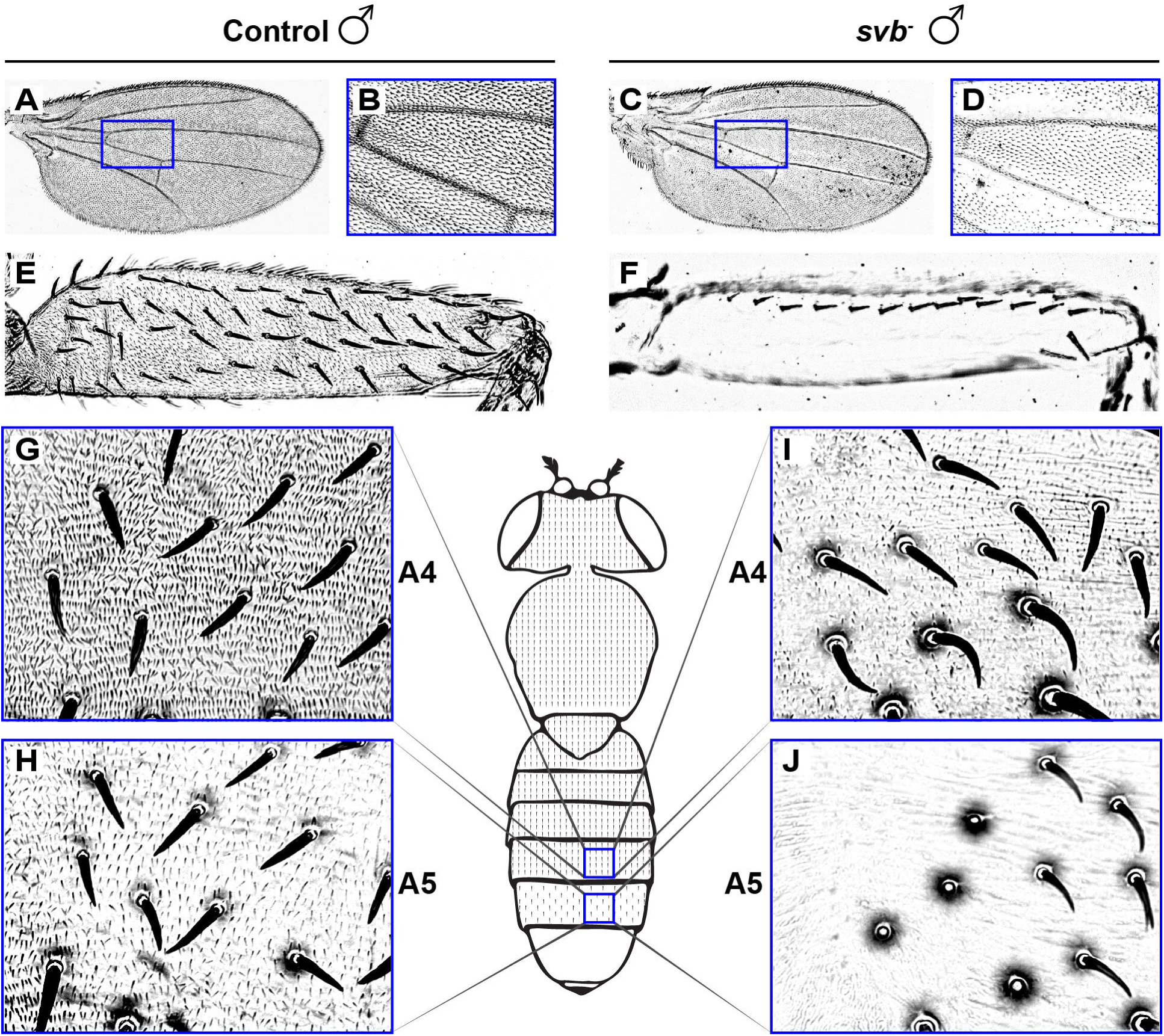
*svb* is required for the production of trichomes in the adult cuticle. (A-J) The cuticle of control f[36a] adult wing (A-B), leg (E) and abdomen (G-H) is covered with trichomes. In *svb* null male escapers (f[36a], svbVY, C-D, F and l-J) many trichomes, but not all, are replaced by naked cuticle. A complete loss of trichomes is observed in legs (F) and abdominal segment A5 (J). Blue boxes within the cartoon demarcate the imaged area.

We observed that no trichomes developed in male *svb* escapers on the dorsal abdominal segment 5 (compare Figs 2H and 2J). This observation stimulated a detailed inspection of the wild type trichome pattern and we found a sexual dimorphism in the shape and size of trichomes in the dorsal abdomen: females produce trichomes of similar density and stoutness on abdominal segments A1 through A5 (SI A Fig), whereas males produce qualitatively different trichomes on abdominal segments A1-A4 vs A5 (SI A Fig). We observed that *svb* expression is lower in abdominal segment 5 versus more anterior segments in both sexes and this difference may contribute to the sexual dimorphism in trichome patterning (SIB Fig).

We confirmed the results observed with the male escapers by generating *svb-/-* clones in the adult. We observed loss of trichomes in *svb-/-* clones in the same regions where trichomes were lost in male escapers (see S2A Fig for an example), confirming that *svb* function is required for the production of some adult trichomes. We also observed loss of trichomes, change in trichome morphology and altered trichome distribution in the head cuticle (S2B Fig). Loss of *svb* function also modified the anatomy of the antennal arista (S2 Fig). In summary, although *svb* is expressed throughout the pupal epidermis (Fig 1H), it is required for the normal development of many, but not all, adult trichomes.

### The embryonic enhancers of svb drive expression in larva and pupa

Since the BAC containing the complete *svb* regulatory region drives expression in embryonic, larval and pupal stages, we wondered whether the previously characterized embryonic enhancers also drove expression in later stages. In the third-instar larvae, we found that of the seven embryonic *svb* enhancers, five drove expression in the epidermis, six drove expression in the foregut and four drove expression in the central nervous system of L3 (Fig 3A and S3 Fig). These results are consistent with the expression of the svbBAC-GFP (Fig 1). In the pupa at 90 h after puparium formation (APF), all embryonic *svb* enhancers drove widespread epidermal expression (Fig 3A-B and S3 Fig.). Most notably, all seven enhancers drive expression throughout the dorsal abdomen (Fig 3B). This level of overlapping expression far exceeds patterns of overlapping embryonic expression that we reported previously [20]. Hence, the seven embryonic *svb* enhancers are both pervasively pleiotropic and redundant.

**Fig 3.**
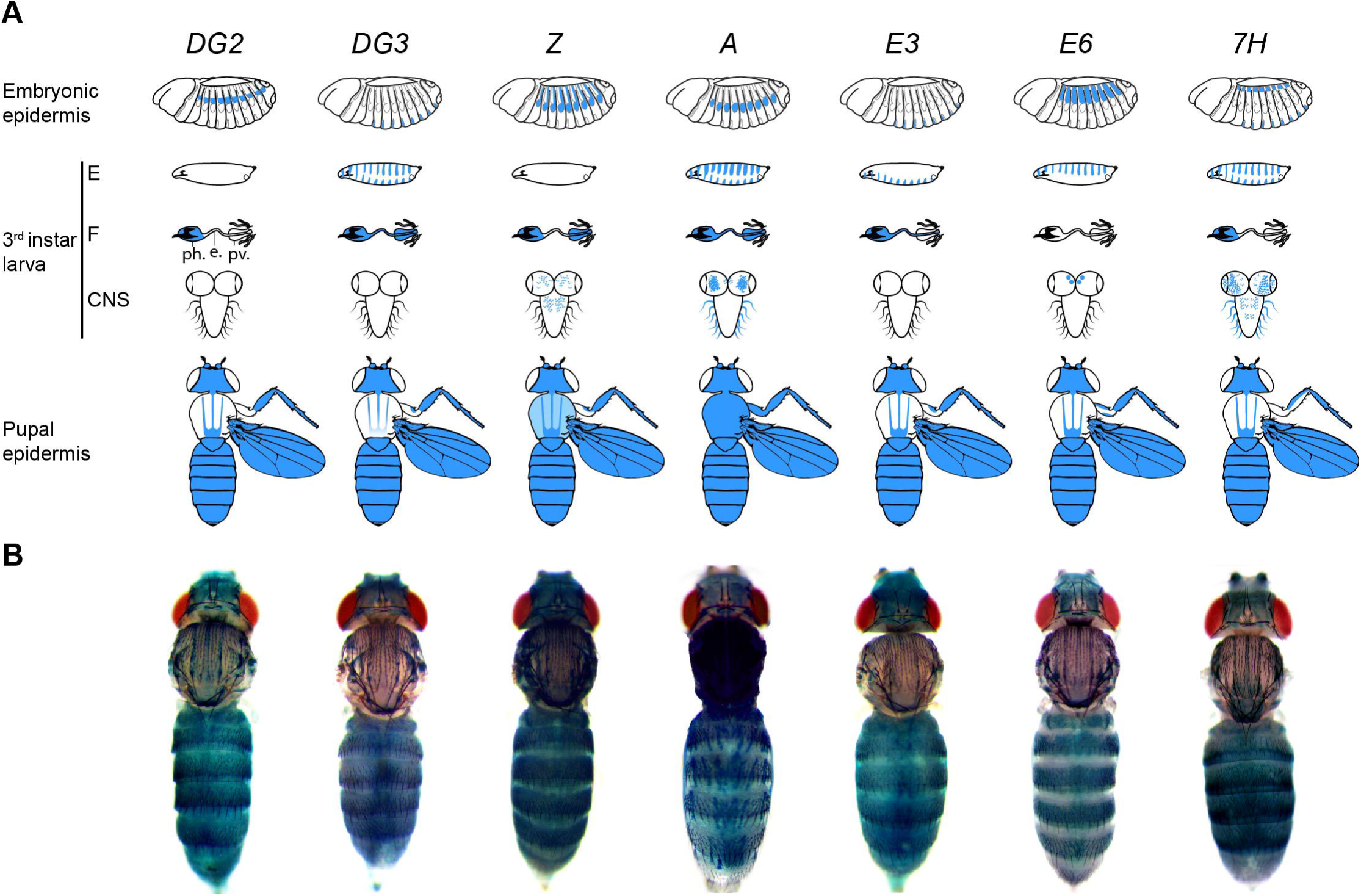
Pleiotropy and redundancy in the activity of the seven *svb* embryonic enhancers. (A) Schematization of the expression pattern driven by each enhancer (blue) in embryo (top), third-instar larva (middle) and pupa (bottom) (see S2 Fig for details). E: Epidermis, F: Foregut, ph: pharynx, e: esophagus, pv: proventriculus, C.N.S: Central Nervous System. (B) Expression pattern generated by each enhancer in the dorsal epidermis of the head, thorax and abdomen (90 hours APF).

### Pupal expression of shavenbaby is conserved in D.sechellia

We have previously shown that five of the *svb* enhancers evolved reduced embryonic activity in *D. sechellia*, a species closely related to *D. melanogaster* [11,20,21]. This loss of enhancer function reduces *svb* expression in *D. sechellia* embryos, causing the loss of many first-instar trichomes [11,17]. In contrast, *D. sechellia* adults, like *D. melanogaster* adults, are completely covered with trichomes (data not shown). To test whether pupal *svb* expression was conserved in *D. sechellia*, we generated a *D. sechellia* svbBAC-GFP with the same genomic boundaries as the *D. melanogaster* svbBAC-GFP (S4A Fig). The *D. sechellia* svbBAC-GFP recapitulated the embryonic expression pattern of *D. sechellia svb*, and no expression was detected in quaternary cells of the dorsal and lateral epidermis (S4B Fig). In contrast, the D. *sechellia* svbBAC-GFP drove GFP expression throughout the dorsal and ventral pupal epidermis, just like the *D. melanogaster* svbBAC-GFP (S4C Fig). Therefore, it is likely that at least some of the D. *sechellia svb* enhancers that lost embryonic expression still drive expression in pupa. To explore this problem further, we examined the embryonic and pupal functions of two evolved *svb* enhancers in more detail.

### The same transcription factor binding sites within E6 are used in both embryo and pupa

We showed previously that the *D. melanogaster E6* enhancer *{melE6)* encodes multiple transcription factor binding sites for the transcriptional activators *Arrowhead {Awn)* and *Pannier [Pnr)* (Fig 4A-B, [17]). *D. sechellia E6 {secE6)* lost four *Awh* sites and acquired a transcription factor binding site for the strong repressor *abrupt*, causing complete loss of its embryonic function (Fig 4C, [17]). We exploited this evolutionary transition to explore pleiotropic roles of *E6.*

**Fig 4.**
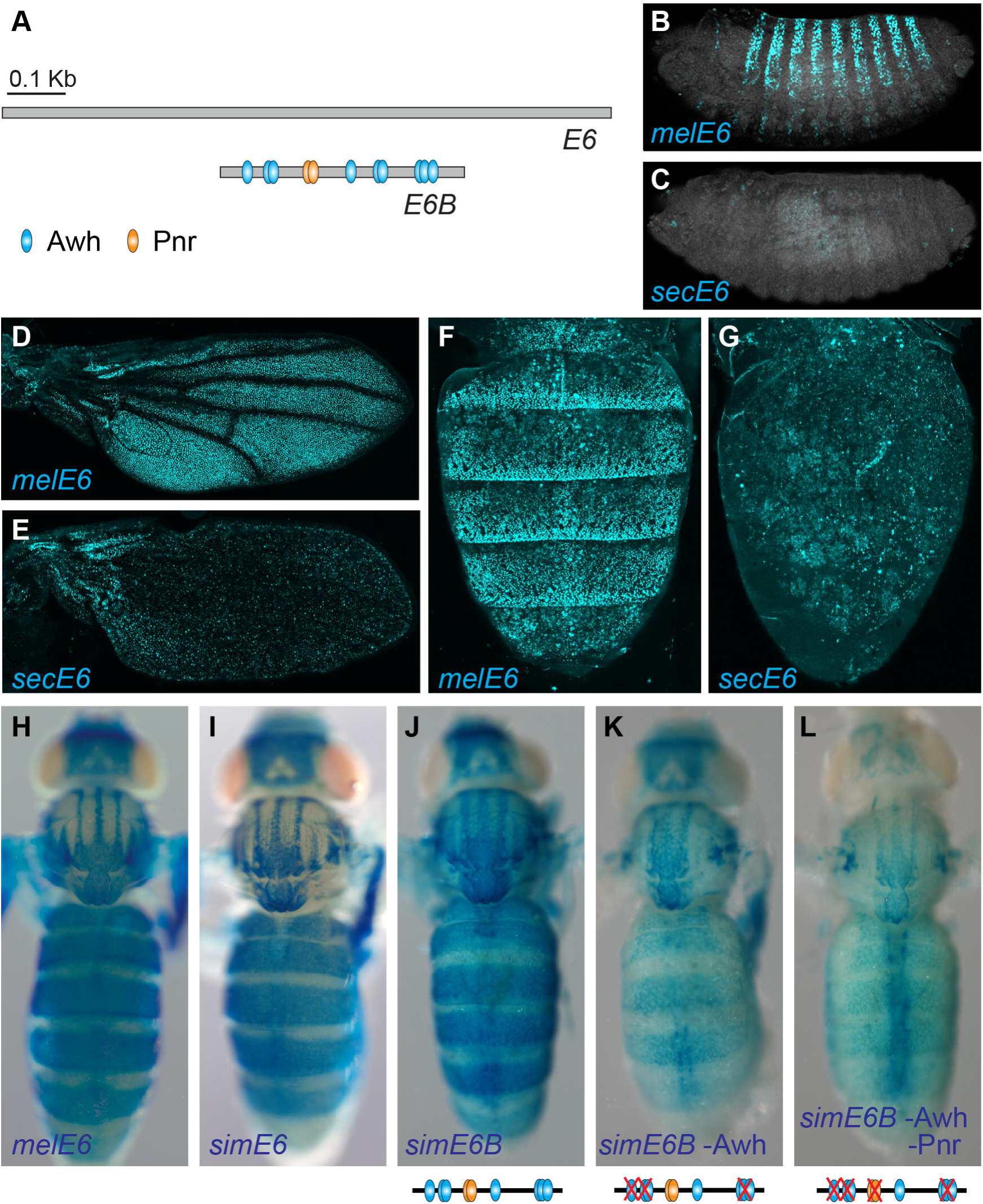
Reuse of TFBSs in the pleiotropic enhancer *E6.* (A) Scheme of D. *melanogaster E6* and *E6B* enhancers. Cyan and orange ovals represent TFBSs for *Awh* and *Pnr*, respectively. (B-C) Expression driven by *D. melanogaster E6* (B, *melE6)* and D. *sechellia E6* (C, *secE6)* in *D. melanogaster* stage 15 embryos. (D-G) Expression driven by *melE6* (D and F) and *secE6* (E and G) in pupal wings (D-E, 74 h APF) and dorsal abdomen (F-G, 84 h APF). (H-L) Expression driven by *melE6* (H), *D. simulans E6* (I, *simE6), D. simulans E6B* (B, J, *simE6B)* and *simE6B* mutants (K and L) in pupa (74 h APF). Red crosses in the schemes below images indicate mutated TFBSs (K, L).

To compare the activities of *melE6* and *secE6* in pupa, we generated new transgenes with fluorescent reporters that were integrated in the same genomic location. We reasoned that if transcription factor binding sites required for embryonic function are required also for pupal expression, then *secE6* should not drive pupal expression. Consistent with our previous observations (Fig 3), the new *melE6* transgene drove strong expression in the wing and dorsal abdomen of the developing pupa (Fig 4D and 4F). In contrast, *secE6* did not drive expression in most of the pupal domains (Fig 4E and 4G). Thus, the evolutionary changes in transcription factor binding sites of *secE6* that led to reduced embryonic expression also reduced pupal expression.

Next, we tested the contributions of individual classes of transcription factor binding sites. We examined a mutated version of the minimal enhancer, *E6B* (Fig 4A), which lacks six *Awh* sites. Like the full length *E6, E6B* drove both embryonic and pupal expression (Figs 4H-J, [17]. Disruption of the *Awh* sites from *E6B* eliminated most pupal abdominal expression (Fig 4K). Additional mutations in the *Pnr* sites led to an even stronger reduction in pupal expression (Fig 4L). All together, these experiments demonstrate that transcription factor binding sites within *E6* are ‘reused’ at multiple developmental stages.

### Different regulatory information generates the various expression patterns of enhancer Z

Next, we analyzed the Z *[melZ)* enhancer, another *svb* enhancer whose embryonic activity was lost in *D. sechellia* [20]. We dissected the ∼4.4 Kb of *melZ* and identified a ∼1.3 Kb minimal enhancer we named *melZ1.3* (S5 Fig). As expected, the orthologous sequence from *D. sechellia {secZ1.3)* did not drive embryonic expression (Fig 5B, [20]). *melZI.3* recapitulated the full Z expression pattern in pupae (Fig 3), including expression in the wings (Fig 5C), legs (Fig 5E), notum and abdomen (Fig 5G). In contrast to our observations for the *E6* enhancer, we found that the *secZ1.3* enhancer drove expression in all tissues where *melZI.3* is active (Figs 5D, 5F and 5H). Therefore, the evolutionary changes that led to the loss of Z expression in D. *sechellia* embryos did not alter Zfunction in pupae.

**Fig 5.**
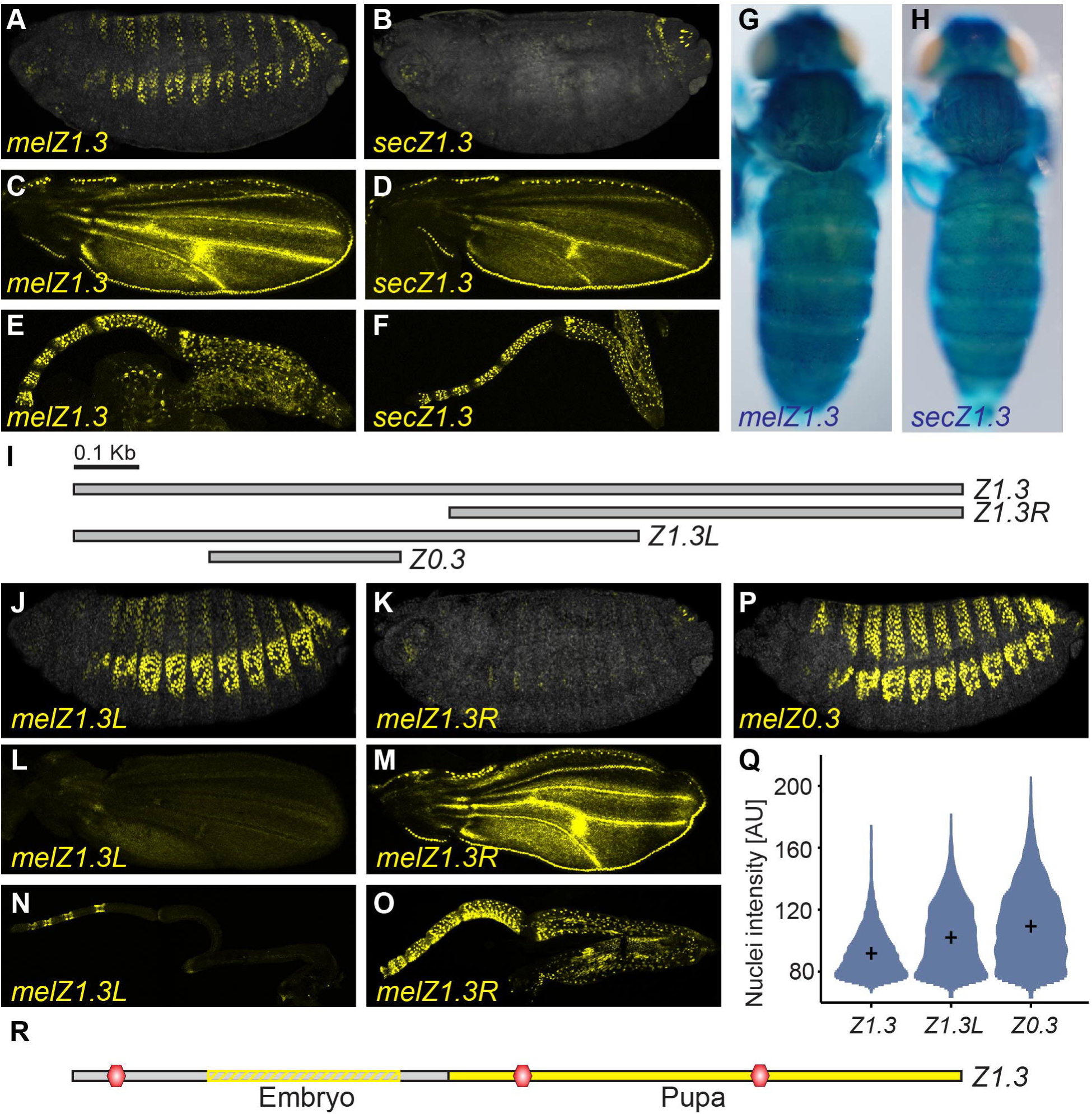
The regulatory information in enhancer Z is partially separated in two modules. (A-B) Expression driven by *D. melanogasterZ1.3* (A, *melZl.3)* and *D. sechelliaZl.3* (B, *secZ1.3)* in *D. melanogaster* stage 15 embryos. (C-H) Expression of *melZI.3* (C, E, G) and *secZ1.3* (D, F, H) in *D. melanogaster* pupal wings (C-D, 36 h AFP), legs (E-F, 36 h APF) and dorsal epidermis of the pupa (G-H, 74 h AFP). (I) Scheme of a subset of the *melZ1.3* enhancer fragments tested with transgenic reporter constructs. (J-P) Expression driven by *D.melanogaster ZI.3L {melZUL), D.melanogaster Zl.3R (melZ1.3R) and D.melanogaster Z0.3 (melZO.3)* in stage 15 embryos (J, K, P), pupal wings (L-M, 36 h AFP) and legs (N-0,36 h AFP). (Q) Quantification of reporter activity in embryos (n=10) carrying *melZ1.3L, melZ1.3R and melZO.3* reporter constructs. Mean intensity is shown with a black cross. (R) Genetic architecture of *melZ1.3. Z0.3* (dashed rectangle) carries most of the regulatory information used for embryonic expression, while Z1.3R (solid rectangle) contains the regulatory information used for pupal expression. Putative binding sites for a transcriptional repressor, which acts in the embryo, are indicated with red hexagons.

To test whether the Z enhancer is divided into discrete modules that drive expression at different developmental stages, we dissected *melZ1.3* into smaller fragments (Fig S1) and tested their ability to drive expression in embryos and pupae. We found that embryonic expression is encoded mostly in *melZUL* (Figs SJ-K), while pupal expression is encoded mainly by *melZIJR* (Figs SL-0 and S5 Fig). In agreement with the results for *secZI.3*, we found that *secZIJR* drove expression in pupal epidermis (data not shown). Thus, the embryonic and pupal expression patterns are encoded by adjacent sequences in the Z enhancer region.

The embryonic and pupal enhancer domains are not strictly separated, however. We identified a 300 bp fragment named *melZ0.3* that drives strong embryonic expression (Fig 5P and S5 Fig). Interestingly, *melZO.3* drove stronger expression than the longer *melZI.3* (Fig 5Q), suggesting that regions outside *melZO.3* contain binding sites for transcriptional repressors, including sites within the pupal enhancer region *melZIJR* (Fig 5R). Nonetheless, the transcription factor binding sites that activate gene expression appear to be present in non-overlapping adjacent domains. Thus, the Z enhancer generates its many functions with different binding sites, which are located in mostly non-overlapping regions.

### Deletion of individual enhancers has contrasting outcomes in embryo and pupa

In recent years it has become evident that the expression of many developmental genes is controlled by multiple enhancers with redundant functions [22]. We have previously demonstrated that the *svb* enhancers drive partially redundant expression patterns in the embryo (Fig 3A), providing phenotypic robustness for larval trichome patterns in the face of environmental and genetic variation [20]. Remarkably, pupae display even greater redundancy of *svb* enhancer expression than embryos (Fig. 3B). We therefore decided to explore the functional consequences of this redundancy.

We used BAC recombineering to generate deletions of individual enhancers (Z7.3, *E6* and *7H)* in the sv6BAC-GFP, and integrated these BACS in a specific *attP* site of the D. *melanogaster* genome (Fig 6A). As an internal control we used a wild-type *svbBAC* with a DsRed reporter (svbBAC-DsRed) that was integrated into a different *attP* site (to avoid transvection effects between BACs). We then quantified expression patterns of the BACs carrying deletions (expressing GFP) relative to the control BAC (expressing DsRed) in the same animal (Fig 6, see Materials and Methods for details).

**Fig 6.**
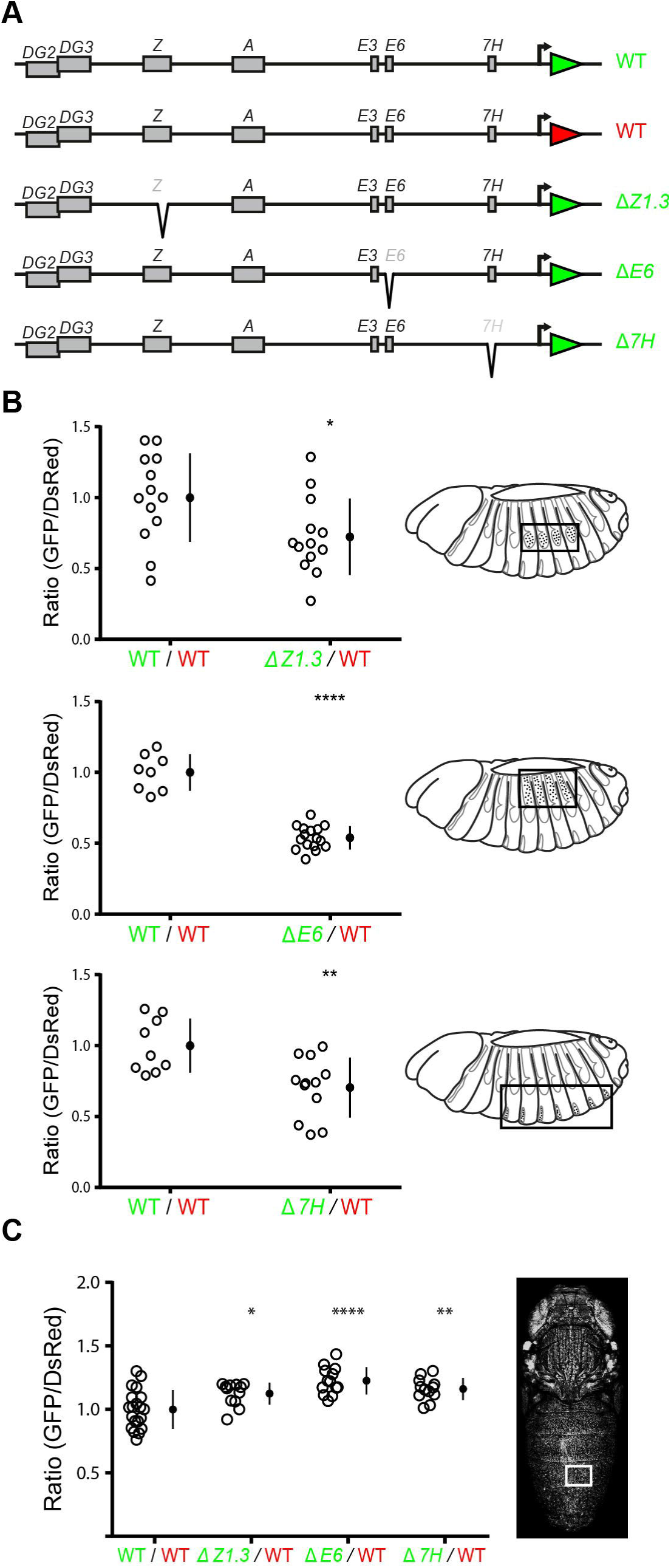
C/s-regulation of *svb* varies between embryo and pupa. (A) Wild type and mutated versions of the svbBAC. The green and red triangles depict the coding sequence of GFP and DsRed, respectively. (B) Effect of enhancer deletions in embryonic expression; ΔZ7.3 (top), ΔE6 (middle) and Δ7H (bottom). Open circles indicate the average ratio (GFP/DsRed) for each individual. Closed black circles and vertical lines indicate mean and one standard deviation, respectively. P values were calculated with two-tailed unpaired t-tests (*p<0.05, **p<0.005, ****p<0.0001). Boxes within embryo cartoons specify analyzed regions. (C) Effect of enhancer deletions on pupal expression. The GFP/DsRed ratio was measured in part of abdominal segment A4 (rectangle) of pupae 90 hours APF. Open circles indicate the average ratio (GFP/DsRed) for each individual. Closed black circles and vertical lines indicate mean and one standard deviation, respectively. P values were calculated with two-tailed unpaired t-tests (*p<0.05, **p<0.005, ****p<0.0001).

Removing *Z1.3, E6*, or 7/7 resulted in a decrease of the mean GFP expression in embryos of 28%, 46%, and 38% respectively (Fig 6B). However, none of the BACs with enhancer deletions drove reduced reporter expression in the pupal epidermis of abdominal segment A4 (Fig 6C). On the contrary, we determined that the deletion of single enhancers slightly increases reporter expression. This fact suggests that the function of *svb* enhancers changes during development.

### Deletion of multiple enhancers in the svb locus does not affect the adult trichome pattern

To examine the importance of this redundancy on the phenotypic output of *svb* function we examined the effects of several deficiencies that remove part of the *svb* regulatory region in the native locus. We used *Df(X)svb^108^*, a deletion of the three most distal enhancers *{DG2, DG3* and Z, [20]) and a larger deletion, *Df(X)svb^106^*, that removes the four most distal enhancers *{DG2, DG3*, Z and *A*, Fig 7A). We showed previously that the *Df(X)svb^108^* line produces a normal number of first-instar larval trichomes when embryos develop at their optimal temperature of 25°C [20]. However, when embryos are grown at a stressful temperature, 32°C, they develop with significantly fewer larval trichomes [20]. We found that *Df(X)svb^106^* produces similar results (data not shown).

**Fig 7.**
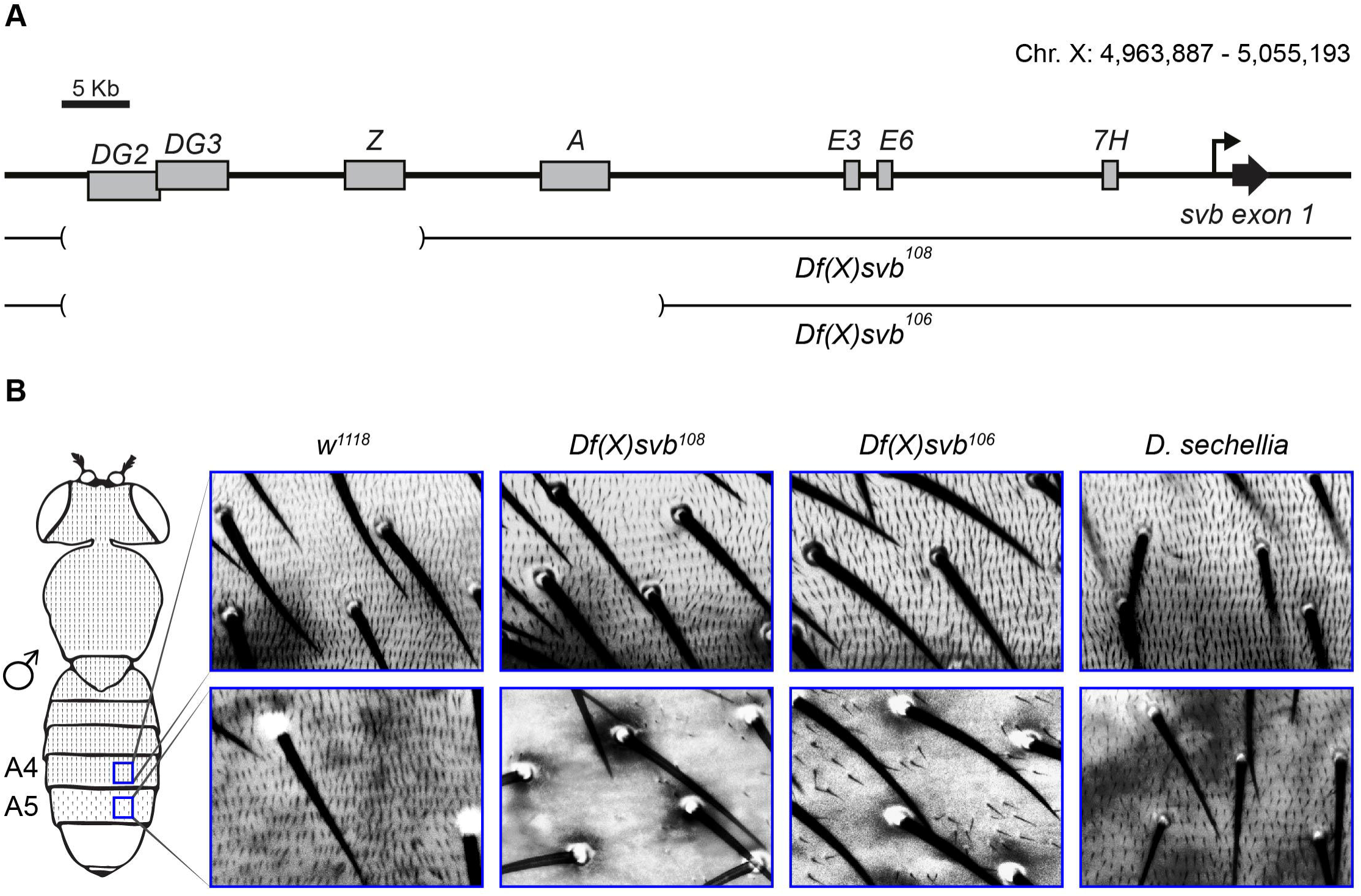
Enhancer deletions in the native *svb* locus alter the A5 trichome pattern only in males. (A) Diagram of the *svb* locus showing the genomic deletions on the X chromosome of lines *Df(X)svb^108^* and *Df(X)svb^m^*. Deletion in *Df(X)svb^m^* removes enhancers *DG2, DG3* and Z, while *Df(X)svb^106^*deletion removes enhancers *DG2*, DG3, Zand *A.* (B) A4 and A5 trichome pattern in adult males of D. *melanogaster, Df(X)svb^108^, Df(X)svb^106^* and D. *sechellia.* Blue boxes within the cartoon demarcate the imaged area.

We could not find any gross changes in the trichome pattern of females (data not shown) when pupae are grown at 25°C. We did notice, however, a small but consistent trichome defect in males carrying either deficiency grown at 25°C: the dorsum of abdominal segment A5 had fewer trichomes than wild type males (Fig 7B). This result is consistent with the phenotype of male *svb* escapers (Fig 2) and the observation that *svb* is expressed in this abdominal segments at lower levels (SI B Fig). Growing *Df(X)svb^108^* or *Df(X)svb^106^* pupae at 32°C did not alter the adult trichome pattern (data not shown). Hence, the adult trichome pattern is largely robust to removal of up to four of the seven *svb* enhancers. In contrast to the effect of these deficiencies on first-instar larvae, stressful growth temperatures in pupa do not significantly alter adult trichome development. Finally, adult *D. sechellia* males displayed an A5 trichome pattern identical to that of *D. melanogaster* (Fig 7B) despite the fact that *secE6* drives reduced expression in the pupal epidermis.

## Discussion

The *svb* gene encodes a master transcription factor that determines the fate of epidermal cells in *Drosophila melanogaster* [14]. It has been known for some time that the embryonic activity of SVB is necessary to pattern the first-instar larva cuticle [13]. In this work we show that *svb* is expressed in several structures of third-instar larvae which have an ectodermal origin (foregut, central nervous system, imaginal discs and epidermis). Furthermore, guided by previous research [14,15], we demonstrate that *svb* is expressed all over the pupal epidermis, and that this expression is required for the development of most of the adult trichomes.

In this work we demonstrate that the seven enhancers of *svb* that drive expression in the embryo also generate expression in third-instar larva and pupa. Thus, the seven enhancers of svb have a pleiotropic role during development. Recently, studies of chromatin conformation showed that the same genomic regions are active in the regulation of gene expression in both developing limb and developing genitalia of mouse [10]. These data strongly suggested that enhancers might be used in entirely different developmental contexts [10]. Later studies reported that the HLEB enhancer of the *Tbx4* gene functions during both limb and genitalia formation in mice, corroborating the idea that the same regulatory element can be active in two dissimilar contexts [23]. Altogether, our data and previous reports [10,23,24] suggest that pleiotropy of enhancer function might be a common feature of c/s-regulatory regions. A possible explanation for the pleiotropic role of enhancers is that these regions of the genome are structurally or topologically special, and that their qualities facilitate the interaction with basal promoters. This idea is supported by the fact that there is conservation in the position of enhancers in distant lineages [25,26]. Alternatively, it can be hypothesized that new expression patterns are easier to evolve within pre-existent regulatory landscapes and, thus, there is nothing special about the position of enhancers in the genome.

There are two fundamentally distinct models by which pleiotropic enhancers could encode expression in different spatio-temporal domains. First, the same transcription factor binding sites could be used to drive expression in different domains or, second, distinct transcription factor binding sites within the same enhancer region could drive different patterns. We studied two pleiotropic *svb* enhancers in detail and found one example that supports each model. For example, we find that Awh and Pnr binding sites, which activate *E6* in the embryo, are also needed to activate *E6* in the pupa. Similarly, it has been shown that Abd-B and STAT binding sites are required for the function of a *Poxn* enhancer in two developmental contexts [27]. In contrast, in the *svb Z1.3* enhancer, we find that transcription factor binding sites that function in the embryo in pupa are mostly independent. This type of architecture might explain how the *Tbx4* enhancer lost its hind-limb function in snakes without losing its activity in genitalia [23]. In summary, we found that there are multiple genetic architectures through which single enhancers can drive pleiotropic expression patterns.

Our results suggest that the classical view of enhancers as strongly modular genomic elements should be reevaluated: transcription factor binding sites can be ‘reused’ during development and enhancer pleiotropy seems to be a common phenomenon. These facts allow us to challenge the notion that conceives enhancers as elements that are active in a single developmental context and evolutionary unconstrained. Often, enhancer function is schematized with a univocal relationship between a DNA fragment and an expression pattern (one enhancer gives only one expression pattern; for example see [28-30]). This schematization, though generally made to illustrate a concept, conveys the wrong idea of enhancers as always being active in a single context. On the other hand, the ‘reuse’ of transcription factor binding sites may impose a limit for enhancers to evolve new functions. Thus, pleiotropic enhancers may sometimes constrain and sometimes facilitate evolution, depending on precisely how pleiotropy is encoded.

We observed extensive redundancy of *svb* enhancer activity in pupal stages. This redundancy far exceeds the redundancy we had characterized previously for the embryonic expression pattern, which is required for phenotypic robustness [17,20]. In fact, we observed that deleting individual enhancers has contrasting outcomes in embryo and pupa. Whereas in embryos the loss of one enhancer diminishes gene expression, in pupa the lack of a single enhancer generates a slight increase in gene expression. Furthermore, flies carrying only three of the total seven *svb* enhancers still produce largely normal adult trichome patterns, even when pupae are grown under stressful conditions. In summary, the function of *svb cis-*regulatory region in pupa appears to result in strong robustness of the adult trichome pattern.

We have shown that *svb* enhancers are pleiotropic and that their expression is highly redundant. Indeed, in *D. sechellia* these enhancers drive enough pupal *svb* expression through stage-specific transcription factor binding sites that the embryonic expression pattern was free to evolve without altering the adult expression pattern. However, it is also possible to imagine a scenario with less redundancy and where pleiotropy is encoded in enhancers through the same transcription factor binding sites (as in the case of enhancer £6), which would strongly constrain the evolution of expression patterns. At present, it is unclear how many enhancers in the genome are pleiotropic, and how their pleiotropy tends to be encoded. Further studies should help determine the extent and encoding of enhancer pleiotropy, clarifying the potential role of enhancer pleiotropy in evolution.

## Materials and Methods

### Genetic constructs and transgenesis

P[acman] CH321-64E24 (bacpacresources.org) contains a 91,307 bp. insert that includes the c/s-regulatory region, the first exon and part of the first intron of *shavenbaby* (www.ncbi.nlm.nih.gov/clone/33521512). We used BAC recombineering [31] to insert a GFP-NLS or a DsRed-NLS in the initiation codon of *svb* to generate sv6BAC-GFP and svdBAC-DsRed and to delete specific enhancers in the context of sv6BAC-GFP. All primers and constructs that were used for BAC recombineering are summarized in SI Table.

The D. *sechellia svb* gene, including the c/s-regulatory region, the first exon and part of the first intron (droSecl: super_4:1,797,878-1,880,229) was subcloned from the BAC DSE1-007L13 (RIKEN BioResource Center DNA Bank) into P[acman]. Subsequently, BAC recombineering was used to insert a GFP-NLS in the initiation codon of *svb* to generate *sec-*svb-BAC-GFP.

The GFP-NLS in pS3AG (a gift from Thomas Williams, addgene plasmid # 31171) was replaced with a DsRed-NLS to generate pS3AR. The DsRed-NLS was released from pRed H-Stinger with enzymes Xhol and Spel. GFP-NLS was removed from pS3AG by cutting with enzymes Xhol and Spel. The pS3AG backbone (without GFP-NLS) was then ligated to the DsRed-NLS. All other transgenes generated in this study were constructed by Genscript (summarized in S2 Table). These constructs were integrated into the fly genome through attP/attB recombination (Rainbow Transgenic flies).

### Fly strains

Enhancer-reporter lines are summarized in S2 Table. In order to generate *Df(X)svb^106^*, pBacPtp4E[f02952] and pBac[f06356] were recombined onto the same X chromosome and a homozygous stock was generated. This stock was crossed to a line containing a hs∷flipase and larvae were heat shocked at 37°C for 1 h each day during larval development. After crossing these adults to *white* flies, we selected adults that had lost one copy of the white-i-transgene (originating on one of the pBac transgenes), which is expected if the two FRT sites recombined to generate a deletion. The deletion was confirmed by PCR, with primers located right outside the deletion (5’- CGTACCGCCTGTTTGCCATA-’3 and 5’-TCCAGACGGAl I I lATGGCC-3), which amplified the expected 7.3 kb fragment containing a pBac transposon. We then generated a stock homozygous for the deletion. *Df(X)svb^108^* was previously described [20]. *D.sechellia* 14021-0248.28 was obtained from the Drosophila Species Stock Center at the University of California.

We generated large clonal territories of *svb’* tissue by employing the Minute technique (Morata & Ripoll, 1975). With this technique clones that contain two wild-type alleles of *Minute^+^* over-proliferate relative to neighboring cells that are heterozygous for a *Minute* null mutation. To mark *svb∼* tissue, we recombined three visible markers (y^1^, w^1^, and f^36a^) onto a chromosome together with a null mutation for *svb* (svb^1^), to generate y^1^ w^1^ svb^1^ f^36a^. We then crossed this strain to flies carrying a dominant Minute allele on the X chromosome. We exposed larvae carrying the genotype y^1^ w^1^ svb^1^ P^6a^ / M to X-Rays (1000 Rad) between 24-72 hours after egg laying. We screened females for clones homozygous for svb^1^ by searching for cuticle containing bristles that were both yellow and forked. We compared trichome patterns in these clones with trichome patterns on flies homozygous for the f^36a^ allele.

### X-gal staining

Third instar larvae were dissected in PBS and fixed in PBS with 4% formaldehyde for 10 min. Staged pupae were removed from the pupal case and then fixed in PBS with 4% formaldehyde for 15 min. After washing in PBT (1X PBS + 0.1% Triton X-100), samples were incubated with X-Gal solution (5 mM K_4_[Fe^+2^(CN)_6_], 5 mM K_3_[Fe^+2^(CN)_6_], 1 mg/ml X-Gal in PBT) at 37°C for 1 hour. The samples were mounted and imaged with bright-field microscopy.

### Immunofluorescence

Stage 15 embryos were collected, fixed, and stained using standard protocols with chicken anti-GFP (1:300, Aves Labs), rabbit anti-RFP (1:150, MBL), mouse anti-pGal (1:500, Promega), anti-chicken AlexaFluor 488 (1:250, ThermoFisher), anti-rabbit AlexaFluor 647 (1:150, ThermoFisher) and anti-mouse AlexaFluor 546 (1:400, ThermoFisher). Pupal tissues were dissected, fixed and stained with mouse anti-fBGal and anti-mouse AlexaFluor 546 as described [32].

### Microscopy and image analysis

Embryos were prepared using standard protocols and immunostained with the antibodies described above. Pupae of the desired stages were removed from the pupal case and placed in a microscope slide for imaging. To analyze the effect of enhancer deletions in svbBACs we measured GFP and DsRed levels in embryos and pupae carrying svbBAC-GFP (WT and deletions) and svbBAC-DsRed (WT). GFP and DsRed signals were measured sequentially over a z-stack in a confocal microscope. Images were analyzed using ImageJ software (rsb.info.nih.gov/ij/). Firstly, background was subtracted using a 50-pixel rolling-ball radius in each slice of the confocal z-stack. Then, we calculated the Sum projection of the z-stacks for each channel in order to compare GFP versus DsRed levels. Max projections were obtained in order to analyze GFP levels between abdominal segments A4 and A5. For embryos, we applied the segmentation masks using the Sum projection of the DsRed channel with the ImageJ autothreshold tool (”IJJsoData dark”). For pupal abdomens, segmentation masks were applied with llastik 1.2.0 software (ilastik.org) to Sum projections of the GFP channel (GFP versus DsRed levels in A4) and Max projections (GFP levels in A4 and A5). We measured the fluorescence mean intensities of each nucleus with the ‘Analyze particles’ tool in ImageJ. Then, we calculated the average of the fluorescence mean intensity of all segmented nuclei. Last, we calculated the ratio GFP/DsRed in each nucleus and calculated the average ratio for all segmented nuclei.

### Cuticle preparation

Adults were collected and frozen until used. Adult cuticles were dissected in PBS and mounted in a microscope slide with a drop of 1:1 Hoyer’s:lactic acid mixture. After overnight drying, the cuticles of adults were imaged with bright-field microscopy. The images were processed using Adobe Photoshop.

## Acknowledgements

We thank Francois Payre for providing the f^36a^ / FM6 stock. We thank Xiaorong Zhang of the Janelia Molecular Biology Shared Resource for help with the sec-svb-BAC-GFP recombineering and the Janelia Fly core facility for help with fly work. We thank the Bioimaging Core Facility at the Technion Rappaport Faculty of Medicine for help with imaging. E.P.B.N is grateful for the generous hospitality of Adi Salzberg and her lab members. E.P.B.N was supported by post-doctoral fellowships from the Human Frontier Science Program. This work was supported in part by Fundacion Bunge y Born and Agencia Nacional de Promocion Cientifica y Tecnologica (PICT 2013-2138) grants to N.F.

## Supporting information

**S1 Fig. svbBAC-GFP expression in pupa and adult trichome patterns**

(A) Adult trichome pattern in abdominal segments A3 to A6 in female (left) and male (right). (B) svbBAC-GFP expression in the dorsum of abdominal segments A2 to A5 of pupae 50 hours APF (left). GFP fluorescence quantification in segments A4 and A5 (right). Each point represents the average mean intensity of all segmented nuclei in A4 and A5 for one individual (n=7). Significance was calculated with a two-tailed paired t-test, *** p<0.0005.

**S2 Fig. *svb* is required for the formation of adult cuticular structures.**

(A-B) Svb” clones on T1 leg (A) and head (B) are outlined in cyan. (C-D) Wild type (C) and subclone modified (D) antennal arista.

**S3 F. Expression driven by the seven *svb* embryonic enhancers in larva and pupa.**

(A-B) X-Gal staining of dissected tissues for each enhancer. (A) Enhancer expression in foregut, central nervous system and epidermis of third-instar larva. (B) Enhancer expression in leg 1 and pupal wing. N.E.: No expression.

**S4 Fig. *shavenbaby [svb)* expression in different developmental stages of *Drosophila sechellia.*** (A) Schematic representation of *D. sechellia svbBAC-GFP {sec-svbBAC-GFP).* Gray boxes represent the seven embryonic enhancers. The site of insertion of the GFP-NLS is indicated in the scheme. (B) *sec-svb-BAC-GFP* expression in stage 15 embryo. Quaternary cells on abdominal segment A2 are outlined. (C) sec-*svbBAC-GFP* expression in dorsal (left) and ventral (right) epidermis of a 90 hour APF pupa.

**S5 Fig. Dissection of the embryonic and pupal functions of *svbZ* enhancer.** (A) Scheme of fragments tested for epidermal enhancer activity with reporter constructs (upper panel). Yellow boxes show fragments with enhancer activity (representative stage 15 embryos are shown in the bottom). Gray boxes indicate no expression. (B) Schematic of a subset of the *svb Z1.3* fragments tested for enhancer activity with reporter constructs. Stage 15 embryos carrying enhancer∷/acZ reporters stained with an antibody against (3-Gal (C) Expression patterns of reporter constructs in pupal dorsal abdomen (74 hours APF) as determined by X-Gal staining.

**S6 Fig. Effect of enhancer deletions on svbBAC-GFP expression.**

(A-B) Representative images of svbBAC-GFP and svbBAC-DsRed expression in abdominal segments of embryo (A) and pupa (B). Black boxes in embryos demarcate analyzed regions for each deletion. The white box in segment A4 of the pupa demarcates the analyzed region.

**S1 Table. List of primers used in this study.**

**S2 Table. List of transgenic lines used in this sudy.**

